# Whole-genome resequencing-based comparative variant analysis identifies candidate genes associated with cross-beak phenotype in Huiyang Bearded chickens

**DOI:** 10.64898/2026.08.11.744104

**Authors:** Fei Ye, Haiyi Yu, Yuyu Hong, Haiquan Zhao, Huiming Kang, Hui Yu, Hua Li

**Affiliations:** School of Life Science and Engineering, Foshan University, Foshan 528225, China; School of Biological Sciences, The University of Western Australia, 35 Stirling Highway, Crawley (Perth), Western Australia 6009 Australia

**Keywords:** Huiyang Bearded chickens, cross-beak deformity, comparative variant analysis, cross-beak severity, *catenin alpha-like 1*

## Abstract

Cross-beaks are deemed a threat to poultry health, productivity, and animal welfare. Nevertheless, due to sporadic cases, heterogeneity of gene loci and incomplete dominance, the molecular mechanism of cross-beak formation, especially the degree of cross, is not yet clear. Thus, we screen key genes and reveal the possible phenotypic formation mechanism of cross-beak by comparison with different degrees of deformity in Huiyang Bearded chickens by compare whole-genome resequencing-based variant analysis. Comparative analysis between cross-beak and normal-beaked chickens identified differential variants in several candidate genes, including *CDH11, CTNNAL1, NRXN3, NRXN1, CDH5, SDC3,* and *DHFR*. Genes harboring these variants were enriched in pathways related to cell adhesion molecules and metabolic processes, with functional annotations involving cell–cell adhesion and neural crest cell migration. Comparative analysis between chickens with severe and slight cross-beak deformities identified additional candidate genes, including *MRPL21, NSUN2, DDX55, GNB3,* and *NFKB2*. These genes were associated with enriched terms and pathways related to focal adhesion, amyotrophic lateral sclerosis, steroid 7 α-hydroxylase activity, and skin-barrier establishment. These findings provide a preliminary catalogue of genetic variants and candidate genes for future functional studies of cross-beak development and severity in chickens.

## BACKGROUND

Cross-beak deformity, also referred to as scissor beak or crossed beak, is characterized by lateral deviation and misalignment of the upper and lower beaks, resulting in incomplete occlusion and impaired beak alignment. Cross-beak deformity has been reported in several avian species and was reported in American brown-headed cowbird, Senegal parrot [1], African seed cracker [2], North American black headed tit [3], Northwest crow [4]. Cross-beak deformity has also been observed in several indigenous Chinese chicken breeds, including Chinese black-boned chickens, bearded chickens, Beijing You chickens and Xiangdong chickens, with reported prevalences of approximately 1%–3% in some populations [5].

Severe cross-beak deformity may impair prehension and feeding, potentially leading to poor growth, reduced survival, and compromised welfare. Nevertheless, some affected birds can survive to sexual maturity, although their reproductive performance and overall production performance may be lower than those of unaffected birds [6, 7].Therefore, elucidating the developmental and genetic basis of cross-beak deformity is important for improving its prevention and management. Genetic factors are considered to contribute to susceptibility to cross-beak deformity [8]. In addition to genetic factors, proposed environmental or developmental contributors include accidental injury, abnormal beak overgrowth, abrasion of the rhamphotheca, exposure to toxins [5, 6], nutritional deficiencies [7], and hypoxia [9]. Phenotypic heterogeneity in cross-beak deformity, together with differences in breed and environmental background, may contribute to variation among studies in the pathways and candidate genes identified.. *Acetyl-CoA acyltransferase 1 (ACAA1)* genes has been proposed as a candidate gene in the development of cross-beak by digital expression spectrum sequencing [10]. *Tudor domain containing 3 (TDRD3), ret proto-oncogene (RET)* and *stathmin 1 (STMN1)* can be candidate genes for beak deformity by genome-wide association analysis and copy-number-variation analysis [7, 11]. A genome-wide DNA methylation study identified *fidgetin-like 1 (FIGNL1)* as a candidate gene potentially involved in mandibular condylar calcification [12]. *Carbonic anhydrase 2 (CA2)* and *carbonic anhydrase 13 (CA13)* have been implicated in the regulation of calcification in the mandibular condyle [13]. In a previous study, we identified bone morphogenetic protein 4 (BMP4) as a candidate gene of potential relevance to cross-beak deformity based on gene-expression patterns and physiological differences observed among facial bones in chickens [14]. Cross-beak deformity is likely a complex trait influenced by multiple genetic and non-genetic factors; therefore, additional studies are needed to prioritize candidate variants and genes and to assess their potential biological relevance.. Whole-genome resequencing enables genome-wide detection of single-nucleotide variants, small insertions and deletions, and, depending on the analytical pipeline and sequencing characteristics, some structural variants in both coding and noncoding regions. Compared with targeted genotyping approaches, whole-genome resequencing can provide broader genomic coverage for exploratory variant discovery. We therefore used whole-genome resequencing to explore variants that may differ among chickens with varying severities of cross-beak deformity..

In this study, we performed an exploratory whole-genome-resequencing-based comparative variant analysis to prioritize candidate variants and genes and reveal the possible phenotypic formation mechanism of cross-beak traits. The findings may provide preliminary targets for validation in larger, independent populations and may ultimately inform future breeding or management strategies aimed at reducing the occurrence of cross-beak deformity..

## MATERIALS AND METHODS

### Animals

All animal procedures were reviewed and approved by the Animal Care and Use Committee of Foshan University (Foshan, China; approval no. FOSU#056). All procedures involving animals were conducted in accordance with the institutional guidelines for the care and use of experimental animals. The animals were from the joint education bases of Foshan University.

A core population of 6,276 Huiyang Bearded chickens was sexed at hatch and managed under standard broiler husbandry and vaccination protocols. A total of 22 chickens with cross-beak deformity from Guangdong Tinoo’s Food Group Co., Ltd. (Guangdong, China) were used as a case group, including eight high (H), seven moderate (M) and seven slight (S) cross-beak chickens, as well as 4 normal individuals (N) as a control group. The classification of high, moderate and slight cross-beak was described previously [14]. Briefly, birds were grouped according to the angle of lateral deviation between the upper and lower beaks: slight, 1 ° –9 ° ; moderate, >10 ° to <19 ° ; and severe, ≥ 20 °. Only peripheral blood samples were collected from all experimental chickens at 105 days of age for genomic DNA extraction.

### DNA preparation

Genomic DNA was extracted from 20 µL peripheral blood using the Whole Blood Genomic DNA Extraction Kit (Aidlab Biotechnologies, Beijing, China) according to the manufacturer’s instructions. Briefly, blood samples were digested with 20 µL proteinase K (20 mg/mL) and 200 µL binding buffer at 70 ° C for 10 min. After cooling to room temperature, 100 µL isopropanol was added, and the mixture was vortexed and transferred to a spin column. The column was centrifuged at 13,000 rpm for 30–60 s and sequentially washed with 500 µL inhibitor removal solution, 700 µL wash buffer, and 500 µL wash buffer, with centrifugation at 12,000 rpm for 30 s after each wash. The column was then centrifuged at 13,000 rpm for 2 min to remove residual wash buffer. DNA was eluted twice with 100 µL elution buffer after incubation at room temperature for 3–5 min and 2 min, respectively. DNA concentration and purity were assessed using a Q5000 microvolume spectrophotometer (Quawell Technology, San Jose, CA, USA). Purified DNA was stored at −20°C until further analysis.

### Library preparation and sequencing

Sequencing libraries were prepared from genomic DNA that met the predefined quality requirements. Genomic DNA was fragmented to an average size of ≤ 350 bp using a Covaris ultrasonicator. Fragmented DNA underwent end repair, during which 3 ′ overhangs were removed by T4 DNA polymerase and 5 ′ overhangs were filled in by DNA polymerase to generate blunt-ended fragments with 5 ′ phosphorylation. The end-repaired fragments were purified using magnetic beads.

Subsequently, an adenine nucleotide was added to the 3 ′ ends of the DNA fragments (A-tailing), enabling ligation of Illumina-compatible paired-end adapters containing complementary 3′ thymine overhangs. Double-stranded sequencing adapters were ligated to both ends of the DNA fragments using T4 DNA ligase. Adapter-ligated DNA fragments of the desired size were recovered and PCR-amplified to generate the sequencing libraries. The amplified libraries were purified before quality assessment. Library fragment-size distribution was assessed using an NGS3K/Caliper system, and the effective library concentration was quantified by quantitative PCR (qPCR). Libraries with an effective concentration of at least 3 nM were considered suitable for sequencing. Sequencing was performed by Novogene Co., Ltd. (Beijing, China) on an Illumina HiSeq X Ten platform using paired-end 150-bp reads (PE150).

### Sequencing Data Quality Control

Raw image data generated by the Illumina platform were processed by base calling to produce raw sequencing reads. Sequencing error-rate distributions and base-quality profiles were examined before downstream analysis. Raw reads were subjected to quality control to generate clean reads according to the following criteria: 1) Paired-end reads containing adapter contamination were removed; 2) If more than 10% of the bases in either read of a paired-end read were undetermined bases (N), the entire read pair was removed; 3) If more than 50% of the bases in either read of a paired-end read had a Phred quality score of ≤5, the entire read pair was removed. The resulting clean reads were used for subsequent alignment and variant-calling analyses.

### Read mapping and single nucleotide polymorphism (SNP) calling

Clean data were mapped to the reference genome (bGalGal1.mat.broiler.GRCg7b) by BWA software (version 0.7.8-r455). Reads with quality lower than Q20 were eliminated during mapping, and BAM files that could be identified and operated by SAMtools software were generated. Based on the mapping results, SAMtools software (version 0.1.19-44428cd) was used to detect SNPs and indels, with the requirement that the number of reads supporting each SNP be no less than 4 and the SNP quality value (MQ) be no less than 20. Then ANNOVAR software (version 2013Aug23) was used to annotate the detected SNPs and indels [15].

Because the number of sequenced chickens was limited, this study was not designed or analyzed as a genome-wide association study. We conducted an exploratory comparative variant analysis to identify variants that differed between phenotypic groups and to prioritize genes for downstream functional interpretation. The autosomal recessive inheritance pattern was employed to conduct SNP screening. To prioritize variants and genes that may be relevant to cross-beak deformity, we identified variants that were shared by chickens in all three affected groups (H, M, and S groups) and absent from the normal-beaked control group. To identify variants potentially associated with deformity severity, To generate hypotheses regarding deformity severity, we identified SNPs detected in the severe group (H) but not detected in the slight group (S), using the same genotype and coverage criteria applied in the primary comparison. Genes harboring these severity-associated differential variants were annotated and prioritized as candidates for further investigation.

### GO annotation and KEGG pathway enrichment analysis

To study the functions of the genes that harbor variants, DAVID 6.8 database tools [8] were used to conduct Gene Ontology (GO) annotation and Kyoto Encyclopedia of Genes and Genomes (KEGG) pathway analysis [18]. The analysis results indicate that a *P* < 0.05 was statistically significant.

### Protein‒protein interaction (PPI) analysis

To further explore potential interactions among proteins encoded by genes harboring candidate variants, we analyzed the PPI through the STRING 11.0 online database. The minimum required interaction score was set to 0.15, and the non-interacting proteins were hidden. Then, the protein‒protein interaction relationship was analyzed by R software (version 4.0.1).

### Sanger sequencing validation

Selected variants were validated by Sanger sequencing. First, blood samples were randomly collected from five chickens with cross-beaks. DNA was extracted according to the instructions of a whole blood genomic extraction kit (Aidlab Biotechnologies, China), as mentioned above. According to the PPI network, *Catenin alpha-like 1 (CTNNAL1), neurexin 3 (NRXN3), and dedicator of cytokinesis 4 (DOCK4)* were selected for Sanger sequencing validation of selected SNP. Primers were designed using Primer Premier 5, and the primer sequences and expected amplicon sizes were listed in Table 1. The PCR products were electrophoresed in 5 μl on a 1% agarose gel, and the target PCR bands were excised from the gel and purified (SanPrep Column DNA Gel Extraction Kit). The PCR products were sequenced with a 3730XL sequencer (ABI, USA).

**Table 1.**
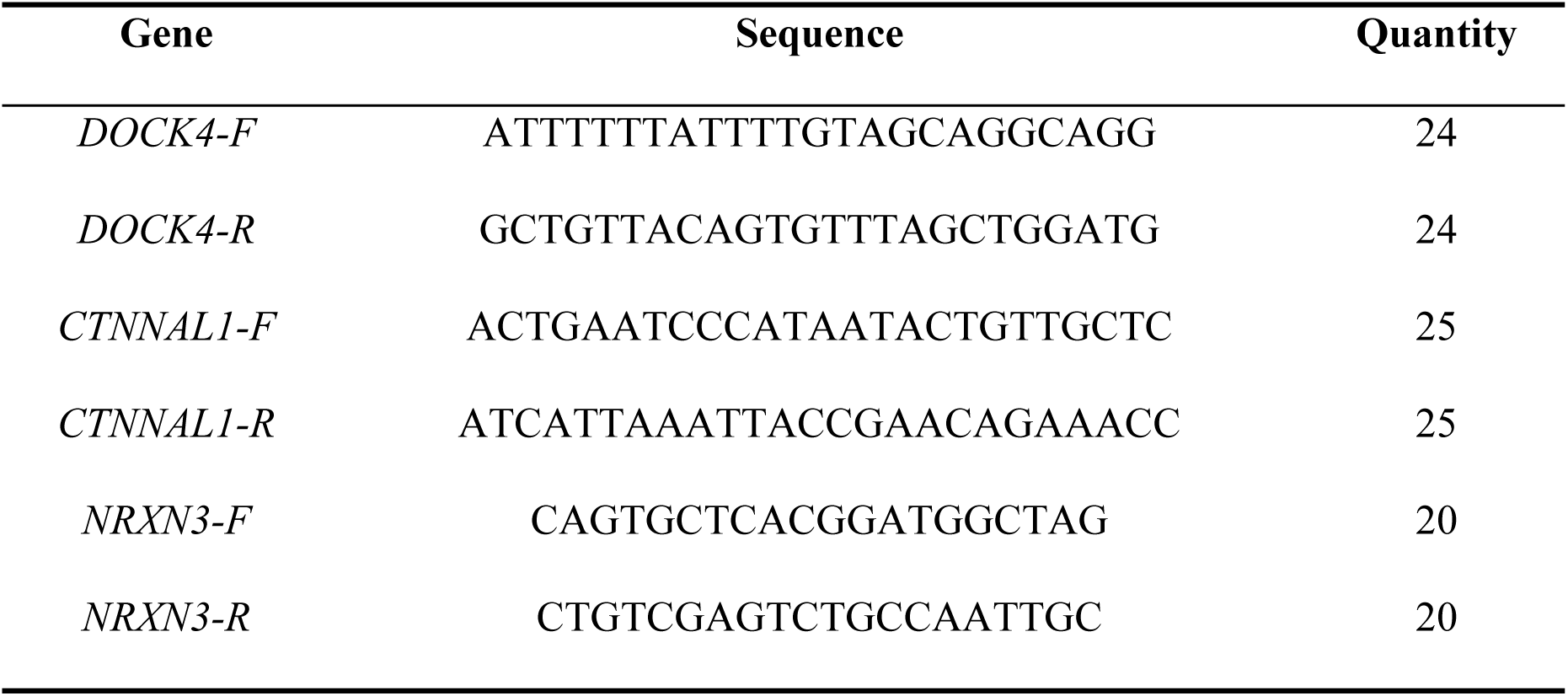
Primer sequences and expected PCR product sizes for Sanger sequencing validation.

## RESULTS

### Whole-genome resequencing quality control

DNA purity was assessed before library preparation, with A260/A230 ratios ranging from 1.8 to 2.4 and A260/A280 ratios ranging from 1.8 to 2.0., indicating that the DNA samples were of acceptable purity for library preparation. The error rate of the intermediate base was less than 0.04%. After the data were processed by quality control, the raw data of each sample were between 12 and 16 G, and the amount of data for all samples was sufficient. The effective rate exceeded 99.43%, among which the bases above Q20 exceeded 95.06%, the bases above Q30 exceeded 89.85%, the base error rate was 0.03%. These metrics indicated that the sequencing data were suitable for downstream alignment and variant-calling analyses. The number of GC bases was between 42.32% and 43.05%, the content and distribution were normal, and the library construction and sequencing were successful (Table 2).

**Table 2.**
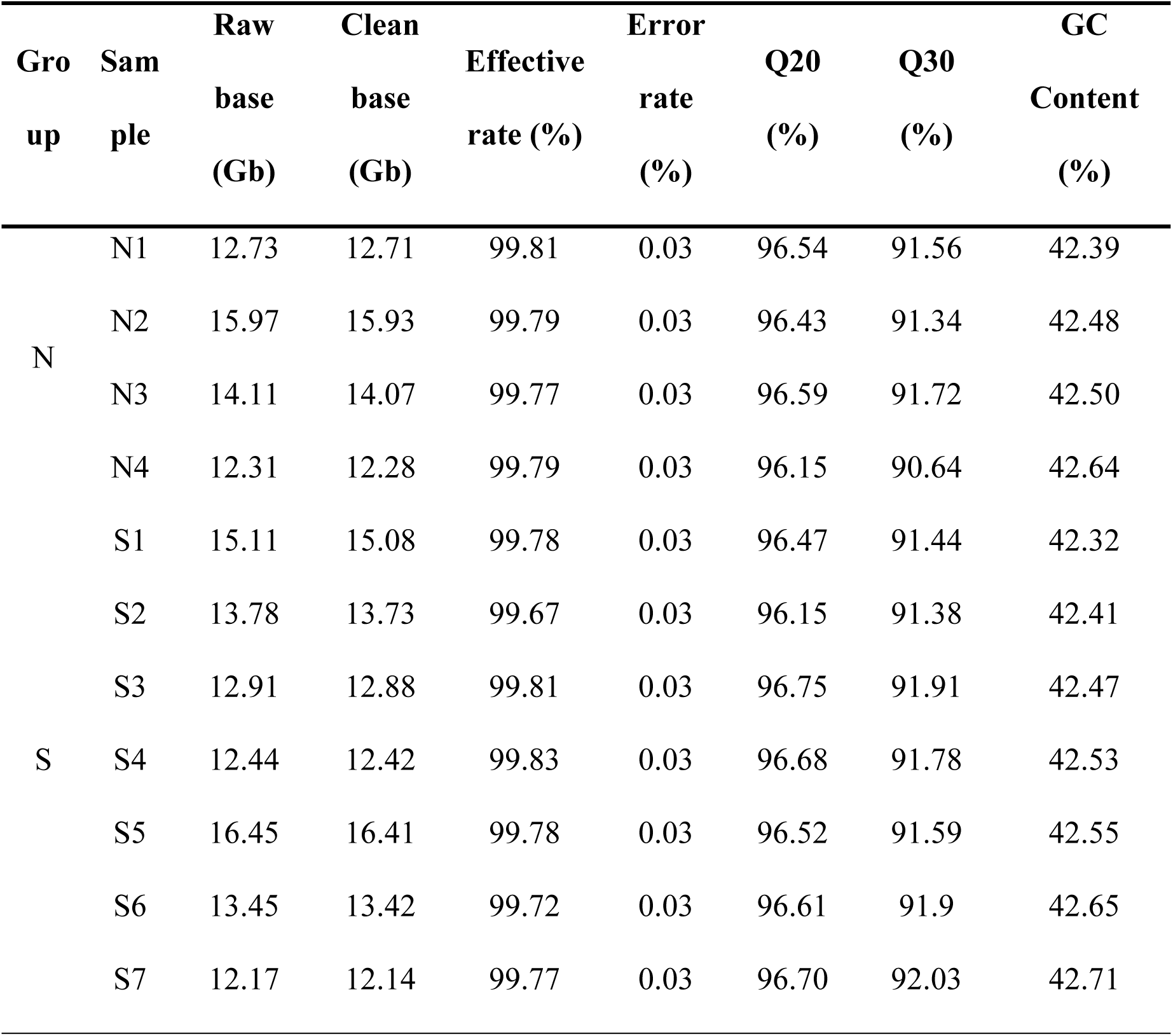
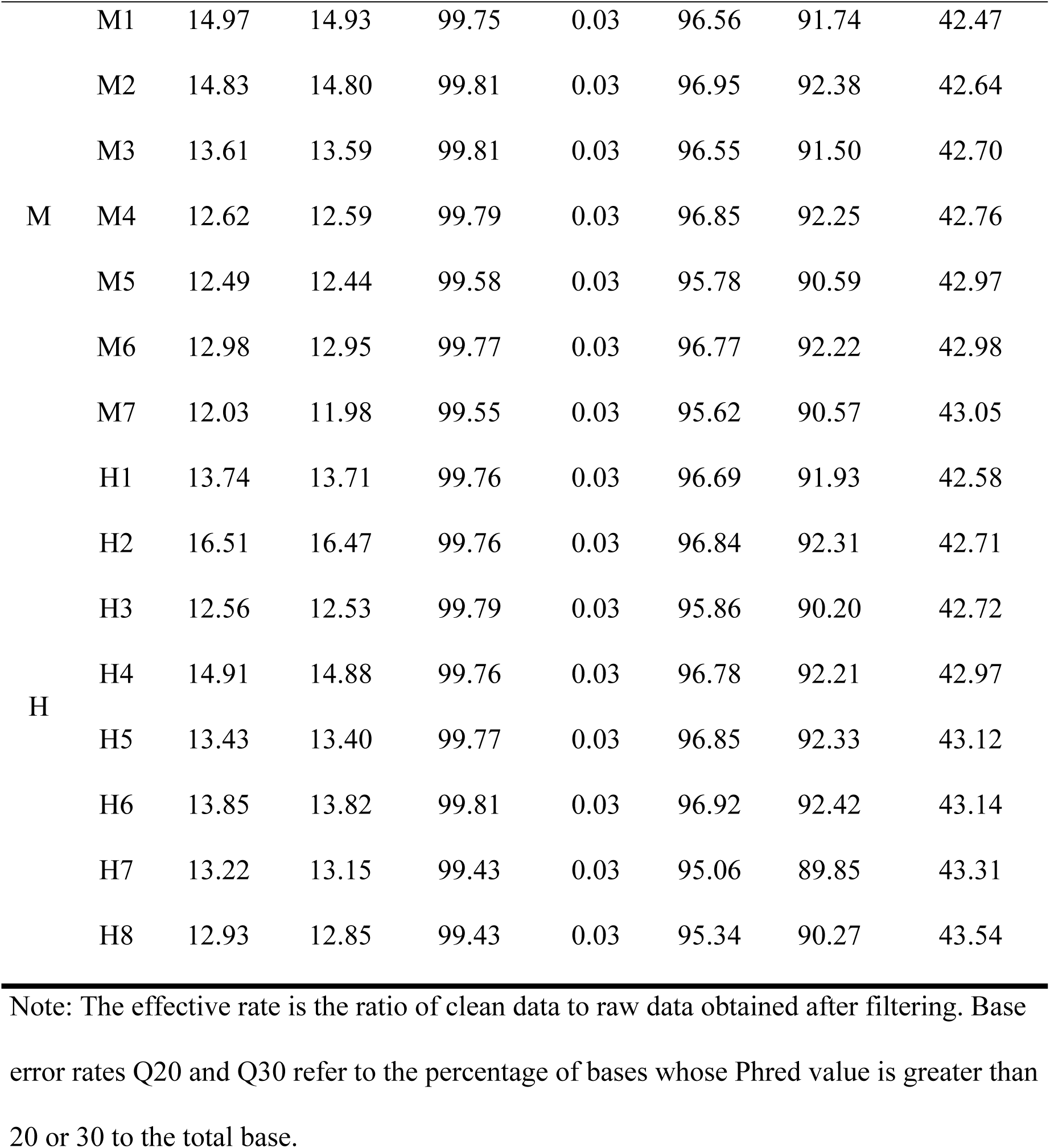
Summary of sequencing data quality.

### SNP detection and annotation

An average of 5.7 M SNP loci were detected and divided into 6 categories in each group (Figure 1). The annotated results were shown in Table 3. The average number of exonic synonymous variants in the normal (N), slight (S), moderate (M), and high (H) groups was 58,484, 59,247, 59,172 and 59,556, respectively. The average number of exonic nonsynonymous variants in the N, S, M, and H groups was 25,151, 25,613, 25,683, and 25,827, respectively. The genome-wide heterozygosity rate (Het rate) of each group was 3.13%, 3.19%, 3.16%, and 3.17%, respectively.

**Figure 1.**
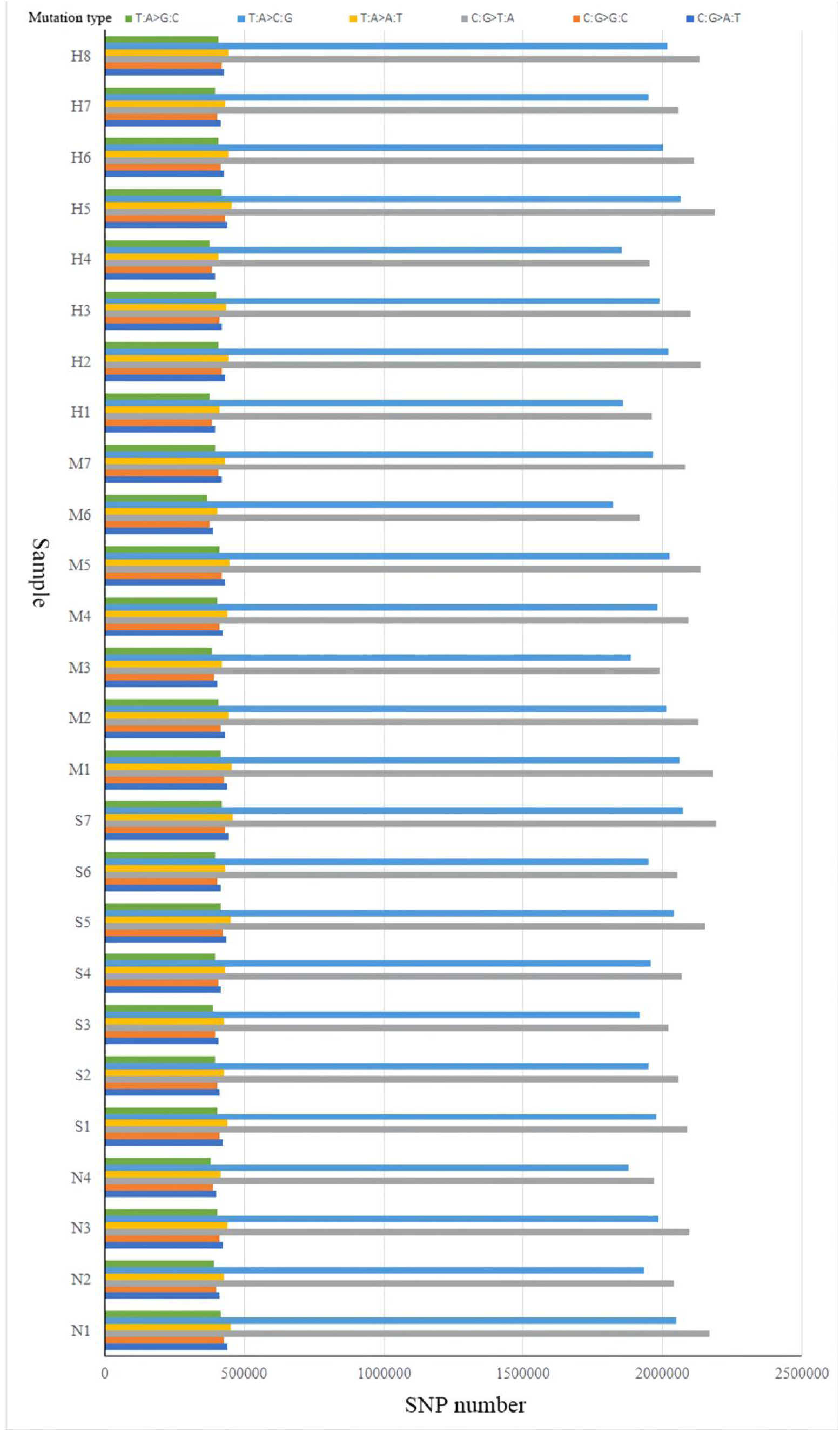
SNP mutation spectrum. Note: The ordinate represents the sample name, the abscissa represents the number of SNPs, and different colors represent different mutation types.

**Table 3.** SNP Annotation Results Statistics.

| Sam<br>ple | Exonic<br>Synony<br>mous | Exonic<br>Non-syn<br>onymou<br>s | Intronic | Upstre<br>am | Downst<br>ream | Intergenic | Het<br>rate<br>(‰) | Total |
| --- | --- | --- | --- | --- | --- | --- | --- | --- |
| N1 | 58,295 | 24,684 | 2,590,736 | 69,995 | 68,111 | 2,729,738 | 3.08 | 5,607,691 |
| N2 | 61,398 | 27,013 | 2,758,051 | 76,165 | 73,837 | 2,884,683 | 3.33 | 5,952,625 |
| N3 | 59,263 | 25,400 | 2,659,682 | 71,090 | 68,613 | 2,812,044 | 3.22 | 5,762,133 |
| N4 | 54,983 | 23,506 | 2,492,633 | 63,984 | 63,395 | 2,672,784 | 2.91 | 5,432,056 |
| S1 | 60,481 | 26,228 | 2,731,206 | 73,367 | 71,233 | 2,890,280 | 3.38 | 5,922,085 |
| S2 | 56,719 | 24,388 | 2,571,193 | 65,456 | 63,819 | 2,717,940 | 3.05 | 5,562,688 |
| S3 | 58,972 | 25,389 | 2,627,061 | 71,040 | 69,684 | 2,724,834 | 3.03 | 5,644,615 |
| S4 | 58,727 | 25,051 | 2,611,128 | 70,595 | 68,579 | 2,748,669 | 3.11 | 5,649,570 |
| S5 | 62,054 | 27,073 | 2,785,568 | 77,144 | 74,380 | 2,917,584 | 3.40 | 6,015,827 |
| S6 | 58,875 | 25,725 | 2,644,468 | 70,308 | 68,425 | 2,806,538 | 3.18 | 5,741,083 |
| S7 | 58,902 | 25,435 | 2,543,837 | 70,773 | 68,922 | 2,771,043 | 3.21 | 5,680,513 |
| M1 | 59,973 | 25,940 | 2,711,360 | 73,000 | 71,650 | 2,830,379 | 3.25 | 5,841,610 |
| M2 | 61,777 | 27,013 | 2,772,576 | 76,324 | 74,143 | 2,896,722 | 3.38 | 5,980,371 |
| M3 | 60,575 | 26,454 | 2,722,214 | 74,987 | 72,283 | 2,848,422 | 3.30 | 5,875,490 |
| M4 | 59,848 | 25,972 | 2,662,691 | 72,790 | 70,904 | 2,789,290 | 3.23 | 5,751,060 |
| M5 | 56,911 | 24,441 | 2,512,614 | 66,512 | 63,945 | 2,683,164 | 3.00 | 5,470,797 |
| M6 | 60,709 | 26,361 | 2,649,790 | 74,206 | 71,592 | 2,749,132 | 3.14 | 5,701,898 |
| M7 | 54,414 | 23,599 | 2,410,188 | 63,972 | 59,940 | 2,602,366 | 2.85 | 5,274,432 |
| H1 | 59,844 | 25,575 | 2,688,424 | 71,986 | 70,162 | 2,825,494 | 3.29 | 5,809,783 |
| H2 | 62,131 | 27,082 | 2,778,074 | 75,737 | 73,202 | 2,912,506 | 3.38 | 5,999,987 |
| H3 | 58,277 | 25,323 | 2,602,518 | 69,865 | 67,786 | 2,764,162 | 3.08 | 5,654,154 |
| H4 | 61,504 | 26,897 | 2,716,773 | 75,567 | 72,317 | 2,836,853 | 3.25 | 5,861,022 |
| H5 | 60,857 | 26,586 | 2,675,809 | 75,366 | 72,667 | 2,780,712 | 3.18 | 5,762,287 |
| H6 | 61,678 | 26,931 | 2,720,945 | 76,613 | 73,485 | 2,819,433 | 3.27 | 5,850,605 |
| H7 | 56,174 | 24,196 | 2,482,325 | 64,748 | 61,616 | 2,638,265 | 2.95 | 5,388,202 |
| H8 | 55,983 | 24,022 | 2,468,717 | 65,705 | 62,114 | 2,638,662 | 2.95 | 5,376,347 |

Note: Upstream and Downstream: The 1-kb regions upstream and downstream of a gene; Exonic: Variants located in the exon region; Syno and Non-syno: Synonymous and missense mutations, respectively; Intronic: Variants located in the intron region; Intergenic: Variants located in the intergenic region; Het rate: Whole-genome heterozygosity ratio, calculated as (number of heterozygous SNPs) / (genome size); Total: Total number of SNP sites.

### SNP and indel analysis of the cross-beak group and normal group

A total of 27 unique nonsynonymous mutation sites in exons of annotated genes were in beak chickens (Table 4). These genes were significantly enriched in positive regulation of synapse maturation, neurotransmitter secretion and cell adhesion (biological process) items, cell surface and integral component of membrane (cellular component) items and calcium channel regulator activity and beta-catenin binding (molecular function) items (Figure 2a and Supplementary file 1). The cell adhesion molecule (CAM) pathway was significantly enriched (Figure 2b and Supplementary file 1). The 27 differential genes were subjected to a variety of protein interaction analyses, and independent proteins without interactions were removed (Figure 2c). The PPI network (Figure 2d) showed that the top ten key genes *cadherin 11 (*CDH11)*, protocadherin 18 (PCDH18), aldehyde dehydrogenase 8 family member A1 (ALDH8A1), cadherin 5 (CDH5), follicle stimulating hormone receptor (FSHR), neurexin 1 (NRXN1), syndecan 3 (SDC3), CTNNAL1, NRXN3,* and *DOCK4* were screened out according to the number of proteins with the same function.

**Table 4.** The unique SNPs annotated genes for the cross-beak group.

| Gene name | Gene symbol | chr | pos | ref | alt |
| --- | --- | --- | --- | --- | --- |
| dedicator of cytokinesis 4 | *DOCK4* | 1 | 27,226,892 | C | T |
| stabilin 2 | *STAB2* | 1 | 54,806,020 | G | A |
| fibronectin type III domain containing 3A | *FNDC3A* | 1 | 170,351,558 | T | G |
| catenin alpha like 1 | *CTNNAL1* | 2 | 88,523,657 | T | C |
| neurexin 1 | *NRXN1* | 3 | 8,022,695 | G | A |
| aldehyde dehydrogenase 8 family member A1 | *ALDH8A1* | 3 | 56,049,130 | T | C |
| zinc finger protein 292 | *ZNF292* | 3 | 76,556,424 | G | A |
| nucleolar protein 10 | *NOL10* | 3 | 96,873,996 | G | A |
| protocadherin 18 | *PCDH18* | 4 | 28,869,545 | A | G |
| neurexin 3 | *NRXN3* | 5 | 40,331,633 | A | G |
| NGFI-A binding protein 1 | *NAB1* | 7 | 54,845 | G | A |
| diacylglycerol kinase delta | *DGKD* | 9 | 1,431,337 | G | A |
| heparan sulfate 6-O-sulfotransferase 1 | *HS6ST1* | 9 | 2,353,708 | T | C |
| leucine rich repeat containing 31 | *LRRC31* | 9 | 20,073,666 | T | C |
| cadherin 5 | *CDH5* | 11 | 11,573,075 | G | A |
| nuclear FMR1 interacting protein 2 | *NUFIP2* | 19 | 6,123,029 | C | T |
| lysosomal protein transmembrane 5 | *LAPTM5* | 23 | 534,447 | T | A |
| nectin cell adhesion molecule 3 | *NECTIN3* | 1 | 89,398,841 | C | T |
| follicle stimulating hormone receptor | *FSHR* | 3 | 80,20,437 | C | T |
| serum/glucocorticoid regulated kinase 1 | *SGK1* | 3 | 56,049,130 | T | C |
| amino adipate aminotransferase | *AADAT* | 4 | 25,168,557 | G | A |
| solute carrier family 7 member 11 | *SLC7A11* | 4 | 28,869,545 | A | G |
| pleckstrin homology domain containing B2 | *PLEKHB2* | 9 | 2,353,708 | T | C |
| cadherin 11 | *CDH11* | 11 | 11,573,075 | G | A |
| MAF bZIP transcription factor B | *MAFB* | 20 | 4,557,756 | G | T |
| syndecan 3 | *SDC3* | 23 | 534,447 | T | A |

note: ref means the reference base, and alt means the variant base.

**Figure 2.**
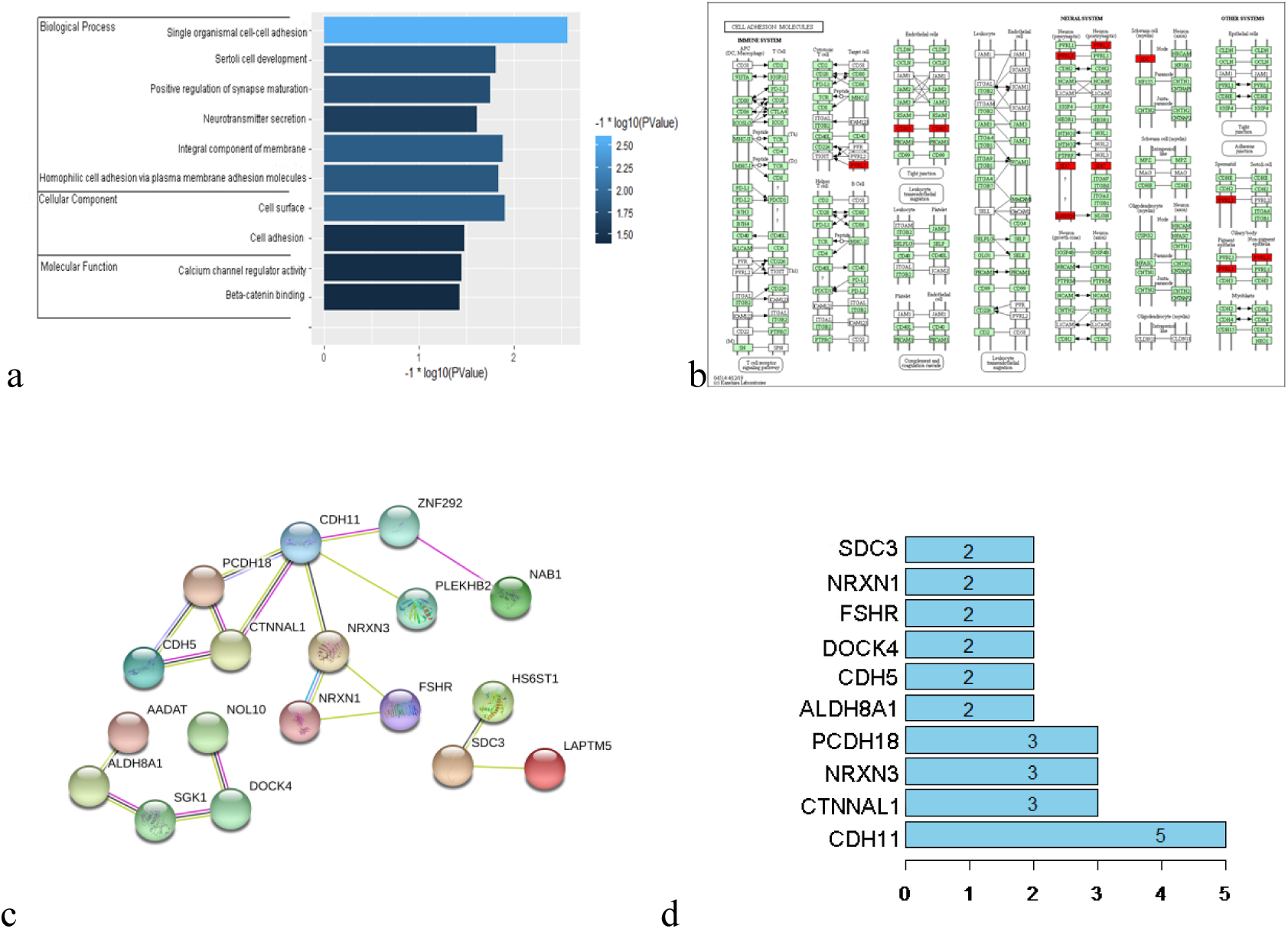
Bioinformatics analysis of SNP variant genes of the cross-beak group and normal group. Note: a, GO enrichment analysis. The ordinate represents the three analysis results of GO: molecular function, cellular component, and biological process; the abscissa represents the -log10 (*P* value) corresponding to each category item, where the lighter the blue color, the more significant the enrichment result. b, KEGG enrichment analysis. Green is marked as a gene specific to the species (gallus), and red is a gene that is unique to cross-beak enriched in the pathway. c, Protein‒protein interaction analysis. d, Key genes of the PPI network. The abscissa represents the number of common functions of genes, and the ordinate represents the abbreviation of genes.

A total of 941 differential genes with unique Indel mutation sites in cross-beak chickens were identified (Table 5 and Supplementary file 2). The biological process most significant term related to cross-beaks was the cellular response to organic substances (Figure 3a and Supplementary file 3). The molecular function most significantly enriched item was the postsynaptic membrane (Figure 3b and Supplementary file 3). The most significantly enriched cell component was cyclo-ligase activity (Figure 3c and Supplementary file 3). In addition, the metabolic pathways were the most significantly enriched (Figure 3d and Supplementary file 4). The differentially expressed genes were analyzed for protein interactions, and the independent proteins were hidden. A total of 323 protein interactions were obtained (Figure 3e and Supplementary file 5). *Dihydrofolate reductase (DHFR)* was deemed the most protein function interaction in the protein network gene (Figure 3f and Supplementary file 6).

**Table 5.** The results of indel annotated genes of the cross-beak group and normal group (partial)

| Gene_name | Gene_symbol | chr | pos1 | pos2 | ref | alt | genotype |
| --- | --- | --- | --- | --- | --- | --- | --- |
| pannexin 2 | *PANX2* | 1 | 2033376<br>1 | 20333<br>761 | - | GTATA | hom |
| galectin 2 | *LGALS2* | 1 | 5121287<br>6 | 51212<br>876 | - | G | hom |
| dehydrogenase E1 and<br>transketolase domain<br>containing 1 | *DHTKD1* | 1 | 6813232 | 68132<br>32 | - | T | hom |
| G2 and S-phase<br>expressed 1 | *GTSE1* | 1 | 1583659<br>4 | 15836<br>594 | - | CAC | hom |
| RNA binding motif<br>protein 17 | *RBM17* | 1 | 4114840 | 41148<br>40 | - | T | hom |
| muskelin 1 | *MKLN1* | 1 | 3740933 | 37409<br>33 | - | CT | hom |
| coiled-coil-helix-coiled-coil-helix domain<br>containing 3 | *CHCHD3* | 1 | 2599752 | 25997<br>54 | T<br>A<br>C | - | hom |
| semaphorin 3A | *SEMA3A* | 1 | 9344407 | 93444<br>07 | - | GCGGC<br>GGGGC | hom |
| semaphorin 3D | *SEMA3D* | 1 | 8840579 | 88405<br>79 | - | GGTGG<br>A | het |
| zinc finger DHHC-type<br>palmitoyltransferase 15 | *ZDHHC15* | 4 | 12,589,3<br>36 | 12,58<br>9,336 | A | - | hom |

note: ref means the reference base, and alt means the variant base.

**Figure 3.**
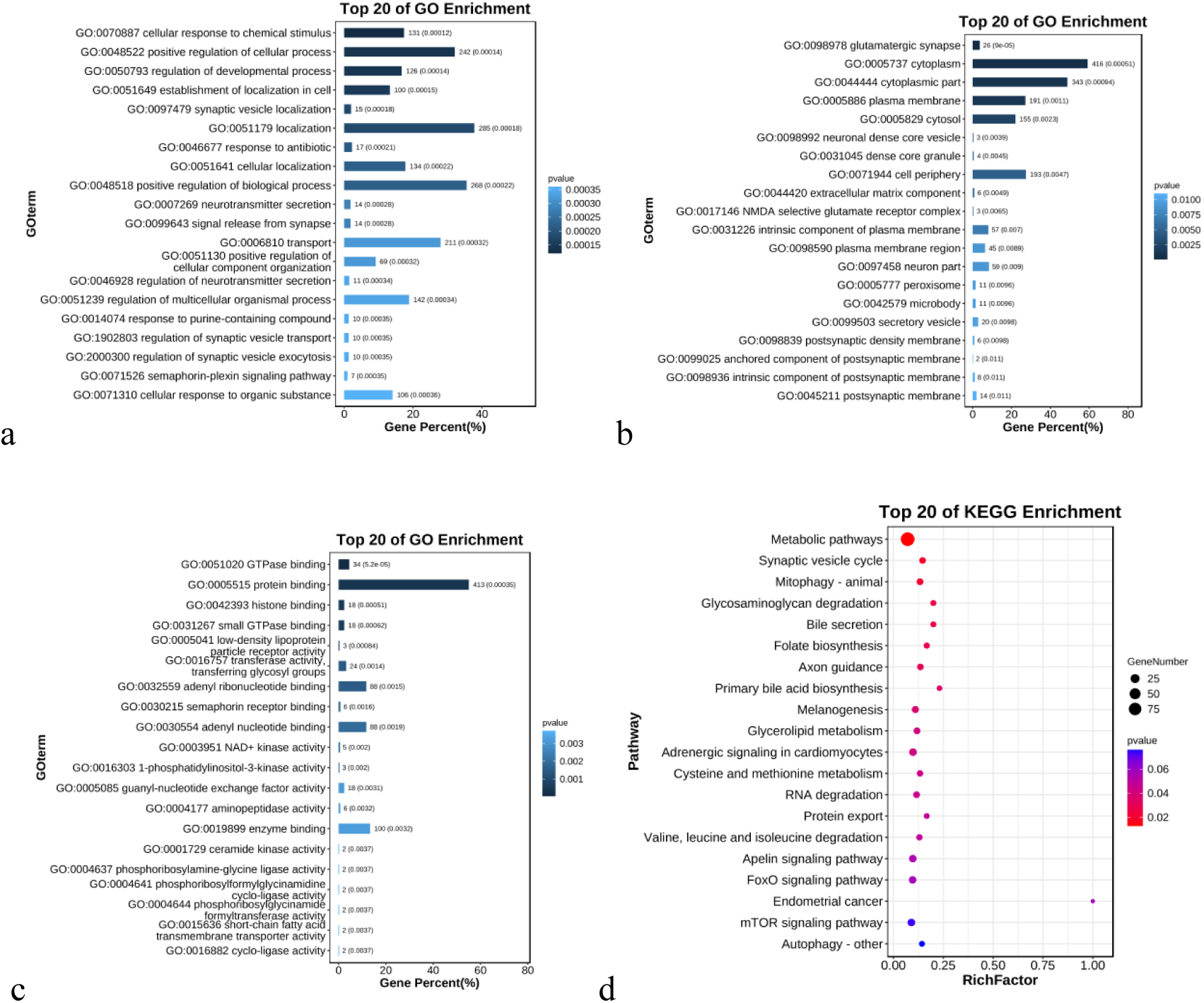
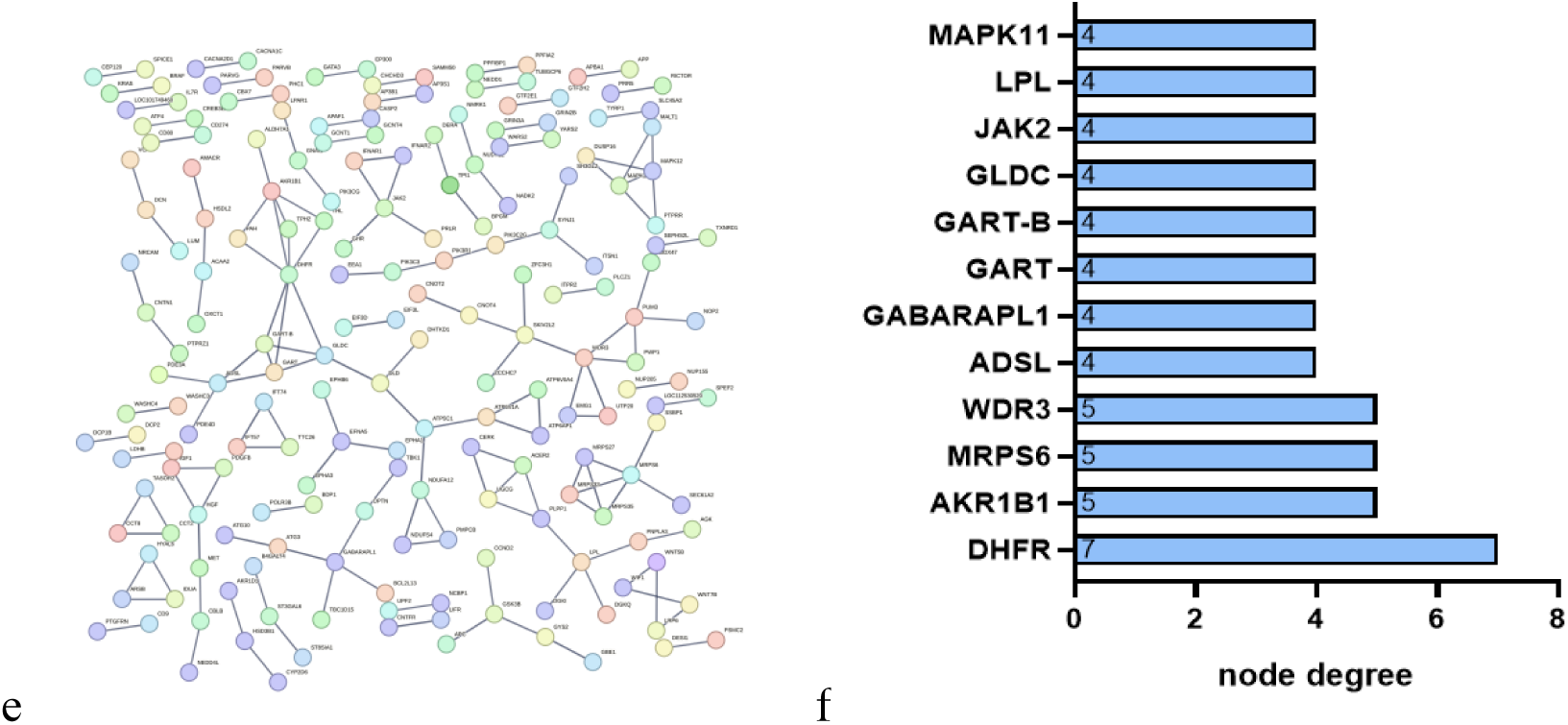
Bioinformatics analysis of indel variant genes of the cross-beak group and normal group. Note: a, Biological process in GO enrichment analysis. b, Cellular components in GO enrichment analysis. c, Molecular function in GO enrichment analysis. d, KEGG enrichment analysis. e, Protein‒protein interaction analysis. f, Key genes of the PPI network. The abscissa represents the number of common functions of genes, and the ordinate represents the abbreviation of genes.

### SNPs and Indels differently analyzed between high and slight cross-beak chickens

A total of 39 differential nonsynonymous mutation SNPs were annotated in genes between high and slight cross-beak chickens (Table 6). These genes were significantly (*P*<0.05) enriched in focal adhesion, amyotrophic lateral sclerosis, and primary bile acid biosynthesis signaling pathways (Figure 4a), which were mainly annotated in steroid 7-alpha-hydroxylase activity of molecular functions (Figure 4b and Supplementary file 7), basal part of cell of cellular components (Figure 4c and Supplementary file 7), and establishment of skin barrier of biological processes (Figure 4d and Supplementary file 7). The protein‒protein interaction analysis results showed that the *mitochondrial ribosomal protein L21 (MRPL21), NOP2/Sun RNA methyltransferase family member 2 (NSUN2), DEAD-box helicase 55 (DDX55)* genes were the core genes of the PPI network (Figure 4e and Figure 4f).

**Table 6.** Different SNPs between high and slight cross-beak chickens.

| Gene name | Gene symbol | chr | pos | Ref | Alt |
| --- | --- | --- | --- | --- | --- |
| ankyrin repeat domain 26 | *ANKRD26* | 1 | 24436260 | A | G |
| MET proto-oncogene, receptor tyrosine kinase | *MET* | 1 | 25100624 | T | C |
| gem nuclear organelle associated protein 8 | *GEMIN8* | 1 | 124164433 | G | A |
| jade family PHD finger 3 | *JADE3* | 1 | 131495616 | T | G |
| collagen type IV alpha 2 chain | *COL4A2* | 1 | 140383611 | G | A |
| spindle and kinetochore associated complex subunit 3 | *SKA3* | 1 | 179935033 | T | C |
| SET domain containing 2 | *SETD2* | 2 | 3989887 | A | G |
| NOP2/Sun RNA methyltransferase family member 2 | *NSUN2* | 2 | 79802912 | A | G |
| anaphase promoting complex subunit 1 | *ANAPC1* | 3 | 3161765 | T | C |
| Ras and Rab interactor 2 | *RIN2* | 3 | 4160854 | A | G |
| aftiphilin | *AFTPH* | 3 | 10379200 | C | A |
| aprataxin and PNKP like factor | *APLF* | 3 | 11975341 | T | C |
| ROS proto-oncogene 1, receptor tyrosine kinase | *ROS1* | 3 | 63137690 | A | T |
| cytochrome P450 family 39 subfamily A member 1 | *CYP39A1* | 3 | 109813086 | T | A, G |
| coagulation factor VIII | *F8* | 4 | 2151744 | T | G |
| centromere protein E | *CENPE* | 4 | 60994801 | A | G |
| mitochondrial ribosomal protein L21 | *MRPL21* | 5 | 1667397 | A | T |
| cytochrome b5 reductase 2 | *CYB5R2* | 5 | 7673369 | C | T |
| cytochrome b5 reductase 2 | *PNLIPRP3* | 6 | 29799424 | A | G |
| family with sequence similarity 171 member B | *ABCA12* | 7 | 4315056 | C | T |
| ATP binding cassette subfamily A member 12 | *ABCA12* | 7 | 4317233 | C | T |
| DEAD-box helicase 59 | *DDX59* | 8 | 1841502 | G | A |
| cytochrome P450, family 2, subfamily J, polypeptide 21 | *CYP2J21* | 8 | 27043561 | A | C |
| solute carrier family 22 member 31 | *SLC22A31* | 11 | 18610623 | G | A |
| DEAD-box helicase 55 | *DDX55* | 15 | 5250966 | A | C |
| SDS3 homolog, SIN3A corepressor complex component | *SUDS3* | 15 | 10102583 | C | T |
| neurofilament heavy polypeptide | *NEFH* | 15 | 11408027 | T | C |
| transmembrane protein 268 | *TMEM268* | 17 | 2890623 | A | G |
| BTB domain containing 17 | *BTBD17* | 18 | 9121883 | C | T |
| calcium activated nucleotidase 1 | *CANT1* | 18 | 9750990 | G | A |
| phosphatidylinositol glycan anchor biosynthesis class S | *PIGS* | 19 | 5883505 | A | G |
| pleckstrin homology like domain family B member 1 | *PHLDB1* | 24 | 5695765 | T | C |
| transmembrane protein 79-like | *TMEM79* | 25 | 2072016 | A | G |
| transmembrane protein 79-like | *LOC107055107* | 25 | 2159932 | C | G |
| SLAM family member 8 | *LZTR1* | 26 | 595342 | C | A |
| leucine zipper like transcription regulator 1 | *LAD1* | 26 | 800470 | A | G |
| paralemmin | *C19orf45* | 28 | 4157290 | G | A |
| TBC1 domain family member 25 | *ZNF541* | 32 | 204768 | G | A |
| tensin 2 | *TNS2* | 33 | 7190238 | G | C |

**Figure 4.**
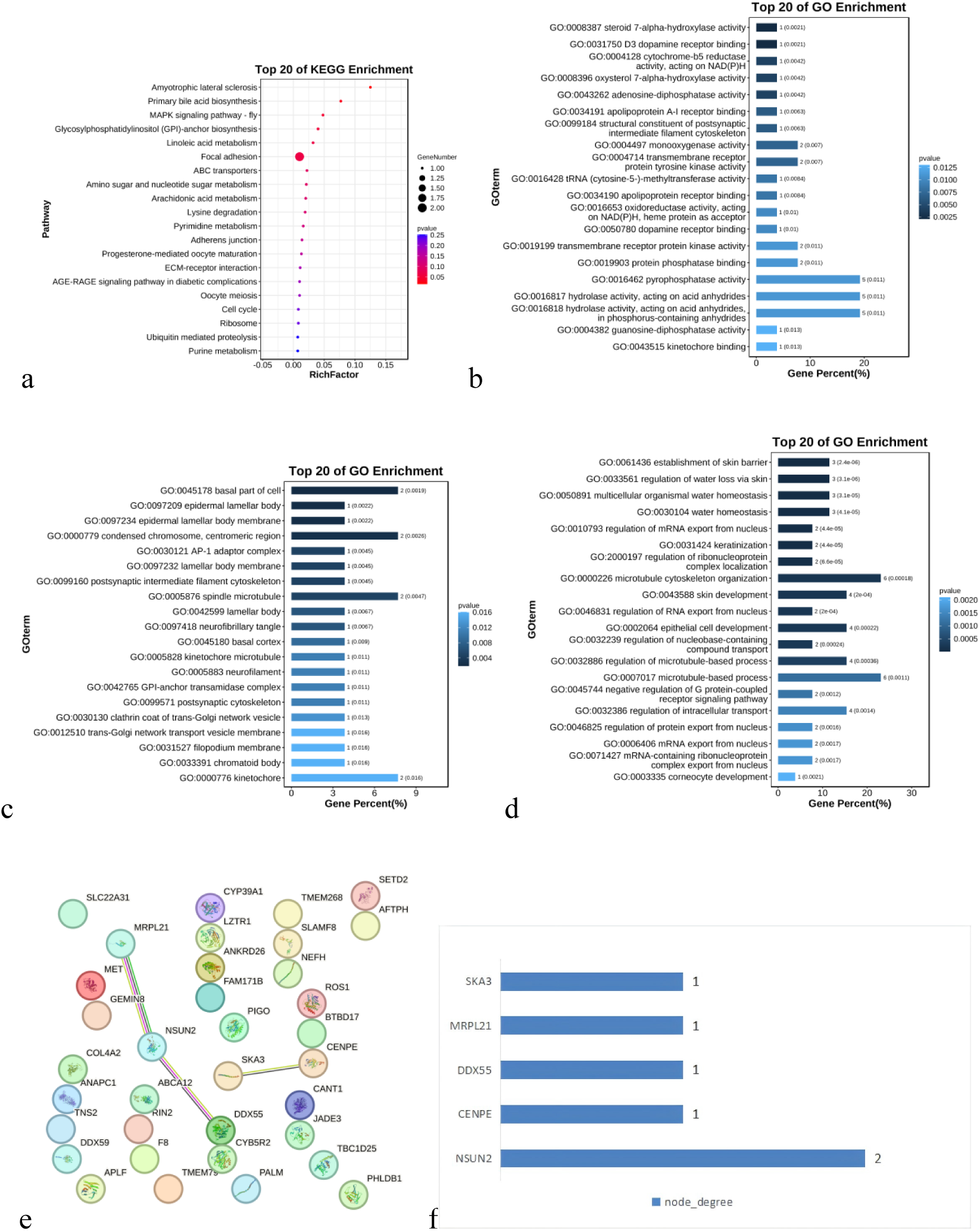
Bioinformatics analysis of SNP variant genes of high and slight cross-beak chickens. Note: a, KEGG pathway analysis; b, GO enrichment analysis of molecular functions; c, GO enrichment analysis of cellular components; d, GO enrichment analysis of biological processes; e, protein‒protein interaction analysis. f, Key genes of the PPI network. The abscissa represents the number of common functions of genes, and the ordinate represents the abbreviation of genes.

A total of 54 differential InDels were annotated in the *G Protein Subunit Beta 3 (GNB3), dedicator of cytokinesis protein 2-like (DOCK2L), and nuclear factor kappa B subunit 2 (NFKB2)* genes (Table 7). These genes were significantly (*P*<0.05) enriched in the chemokine signaling pathway, c-type lectin receptor signaling pathway, relaxin signaling pathway, parkinson’s disease, morphine addiction and dopaminergic synapse pathways (Figure 5a and Supplementary file 8), which were mainly associated with leukotriene-A4 hydrolase activity and interleukin-12 receptor binding of molecular function (Figure 5b and Supplementary file 9); perinuclear theca and I-kappaB/NF-kappaB complex of cellular component (Figure 5c and Supplementary file 9); and cellular response to cholesterol and cellular response to sterol of biological processes (Figure 5d and Supplementary file 9). There were interactions between *signal peptidase complex subunit 1 homolog (SPCS1)* and *TAR (HIV-1) RNA binding protein 1 (TARBP1)* genes (Figure 5e and Figure 5f).

**Figure 5.**
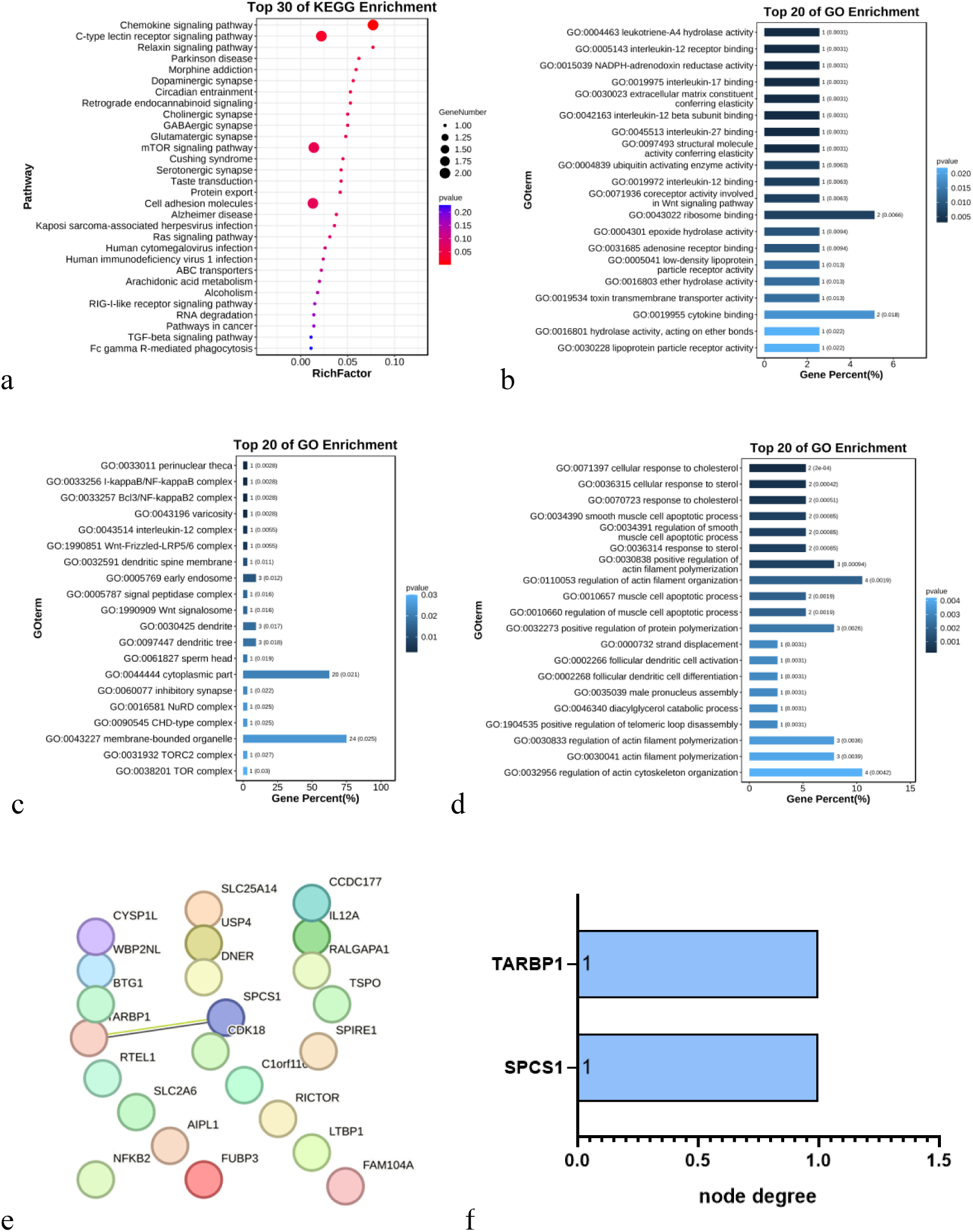
Bioinformatics analysis of indel variant genes of high and slight cross-beak chickens. Note: a, KEGG pathway analysis; b, GO enrichment analysis of molecular functions; c, GO enrichment analysis of cellular components; d, GO enrichment analysis of biological processes; e, protein‒protein interaction analysis. f, Key genes of the PPI network. The abscissa represents the number of common functions of genes, and the ordinate represents the abbreviation of genes.

**Table 7.**
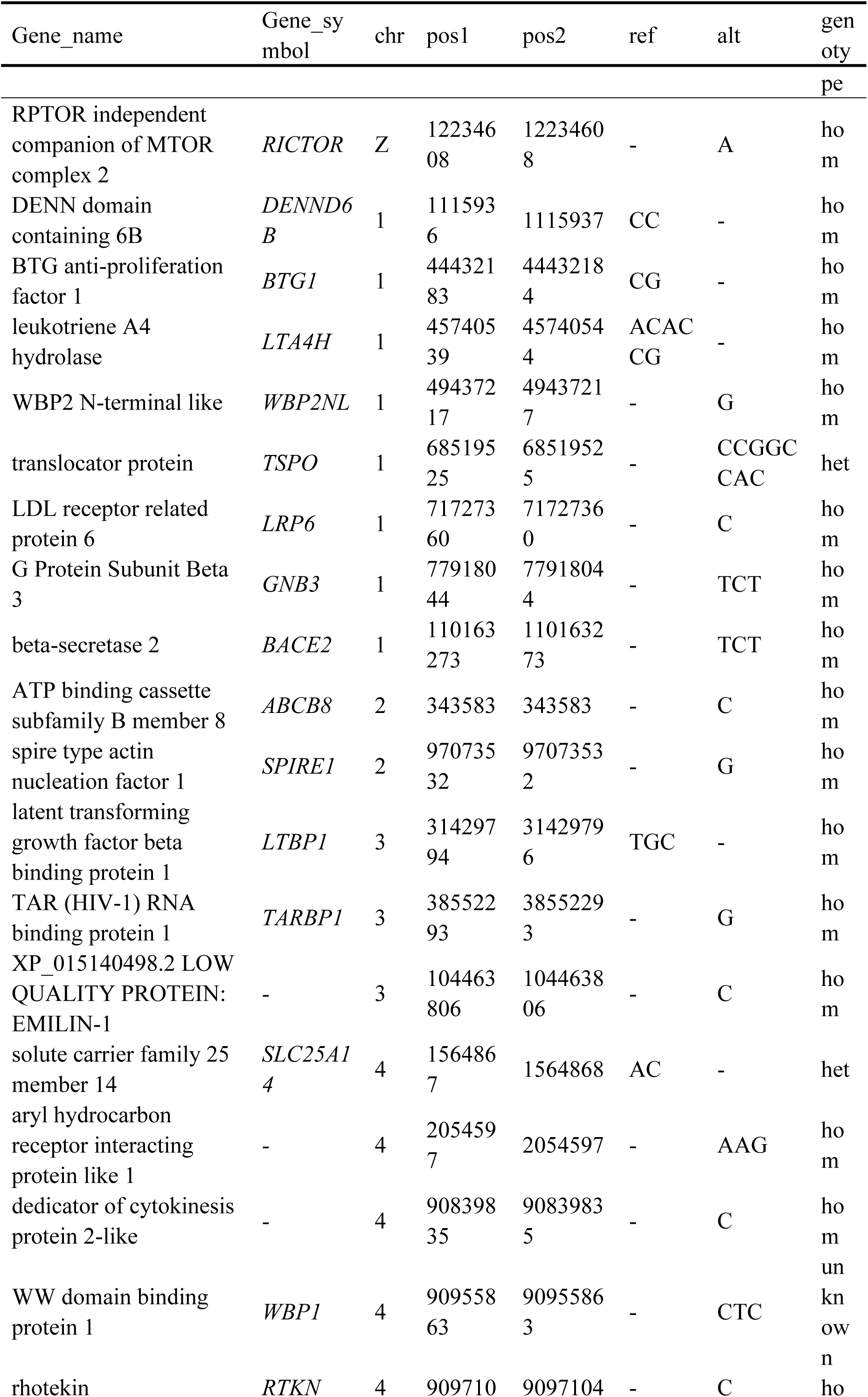
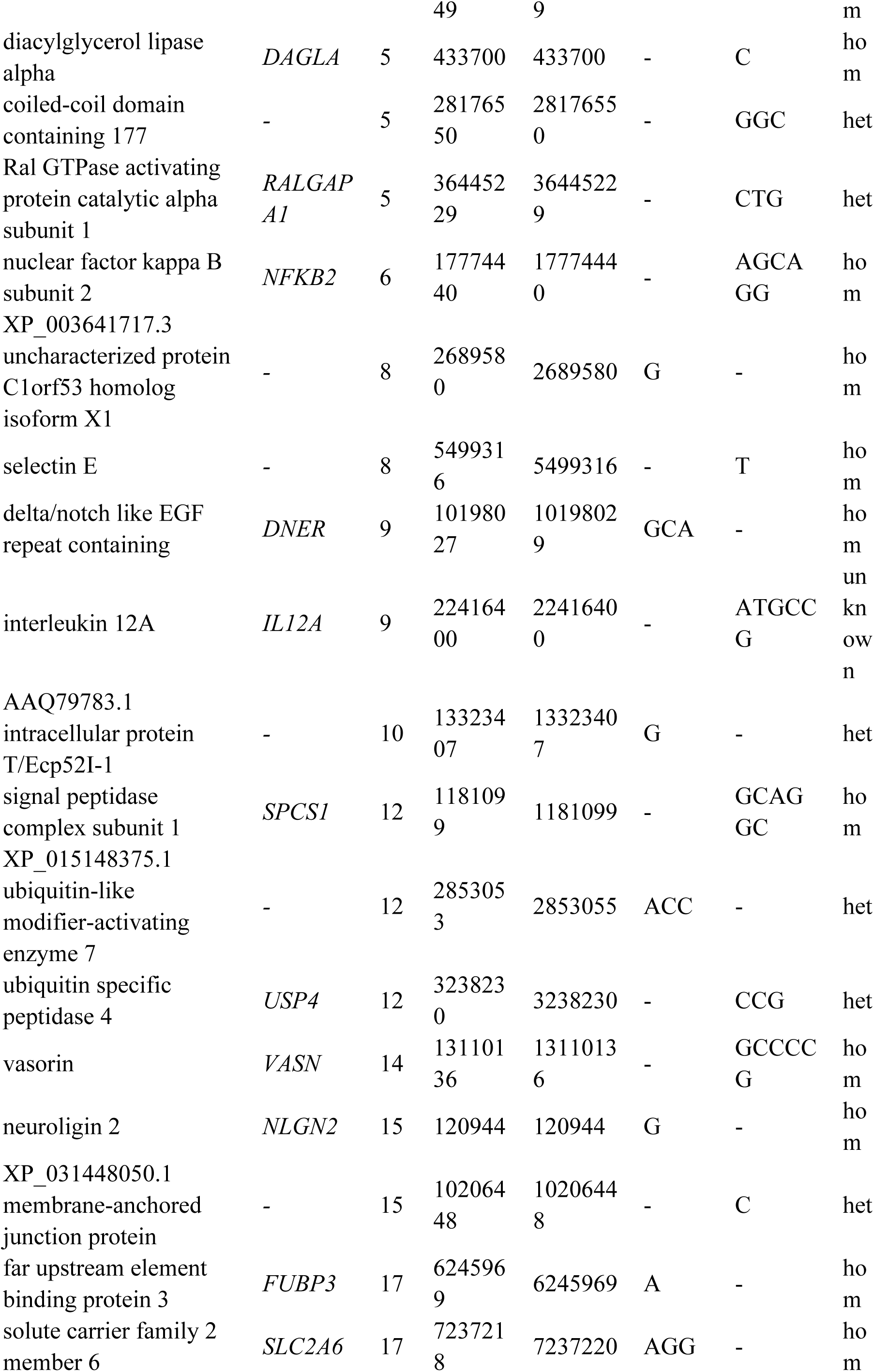
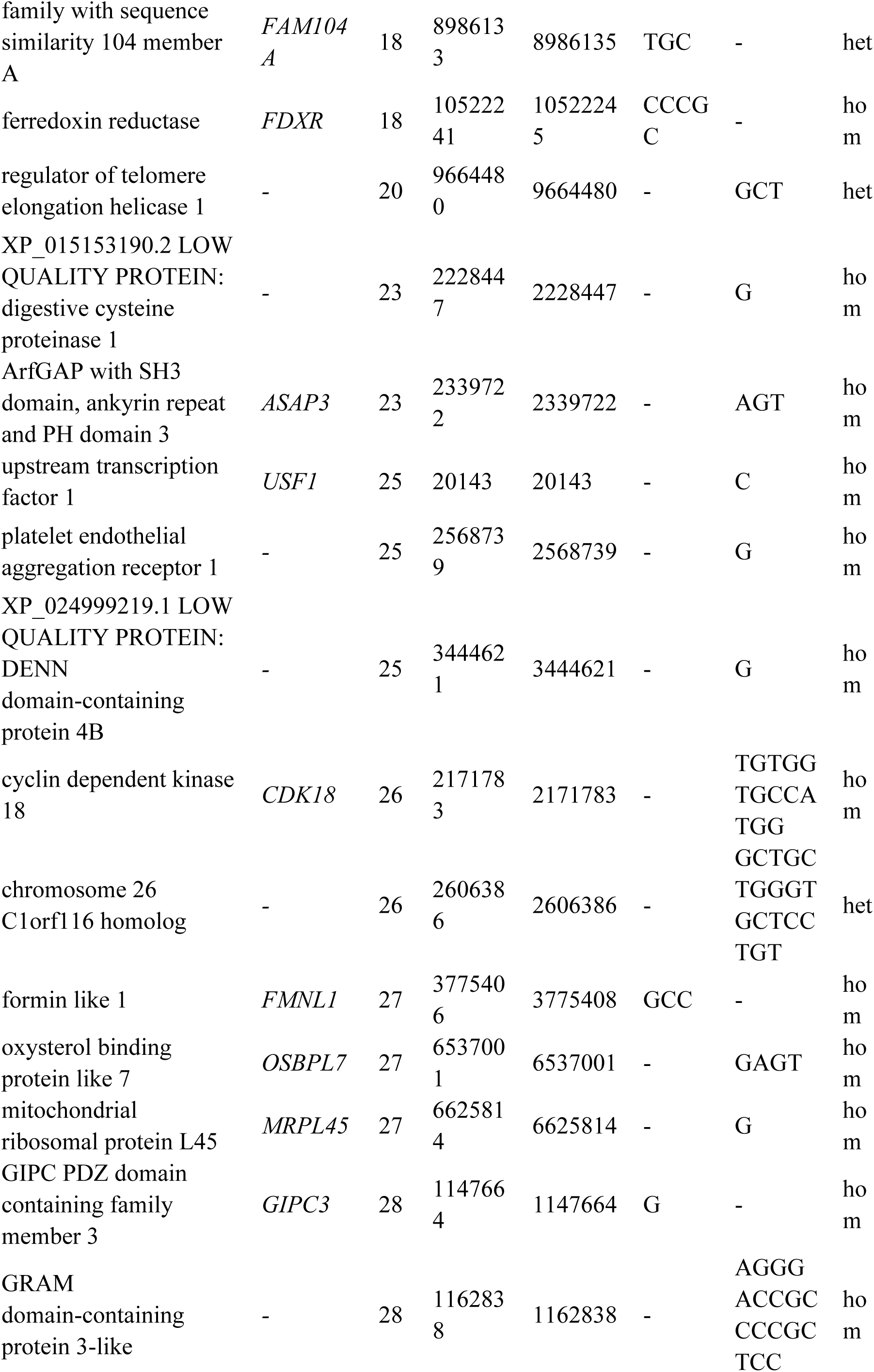
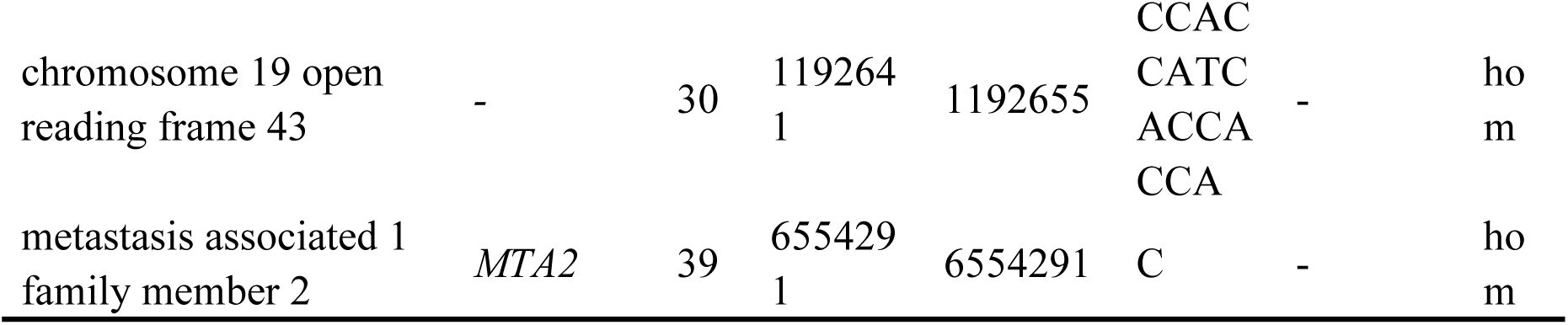
Different Indels between high and slight cross-beak chickens.

### Sanger confirmation of resequencing

The Sanger traces for the *DOCK4, CTNNAL1, and NRXN3* genotypes are shown in Figure 6. Sanger sequencing of the *CTNNAL1* genotype Chr2:88523657 T>C and NRXN3 genotype Chr5: 40331633 A>G were matched with whole-genome resequencing.

**Figure 6.**
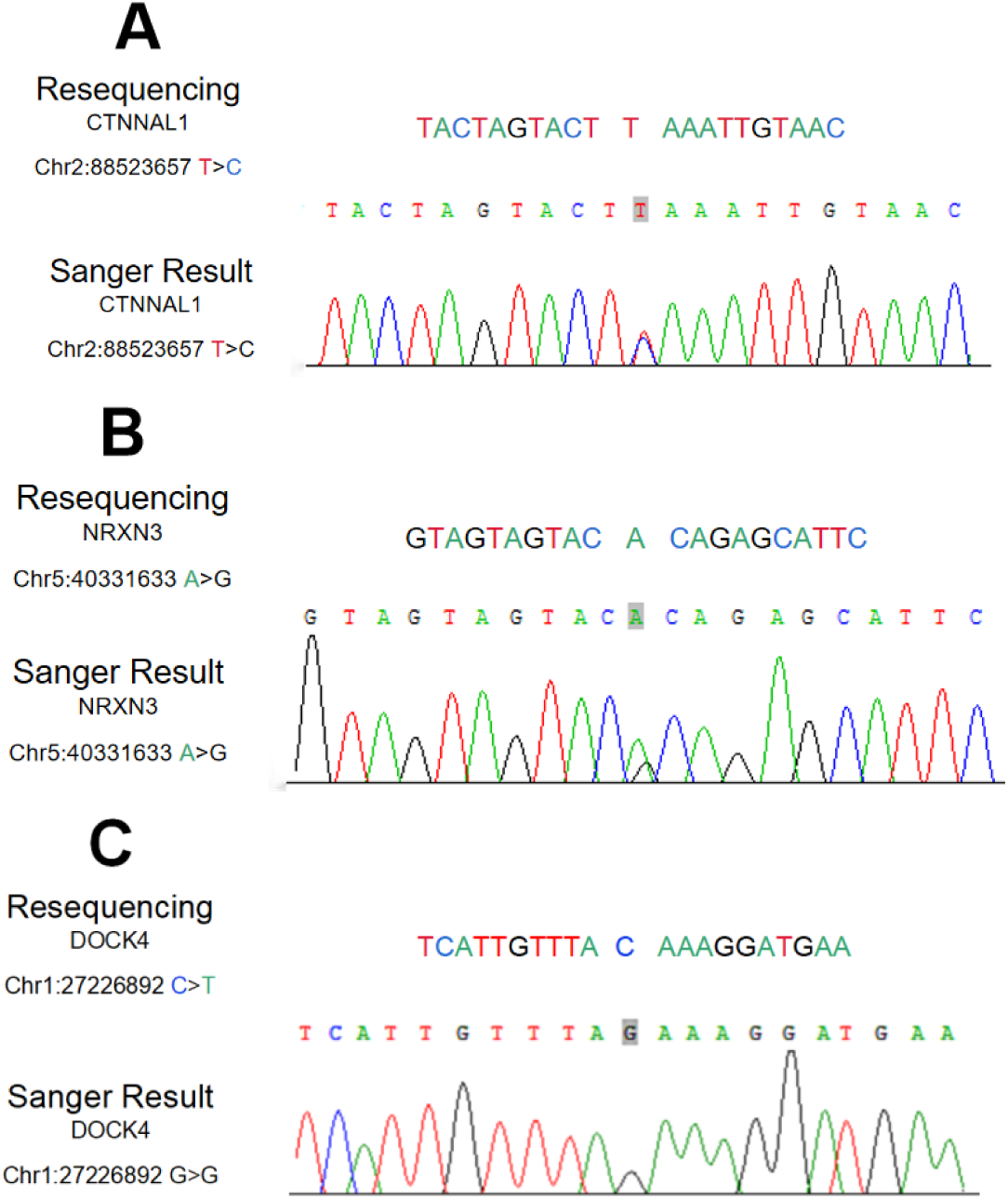
Gene resequencing variant confirmation using Sanger sequencing.

## DISCUSSION

Whole-genome resequencing provides a genome-wide resource for characterizing genetic variation, including SNPs and indels in coding and noncoding regions. In the present study, we used exploratory comparative variant analysis to identify variants that differed between normal-beaked and cross-beak chickens and between chickens with severe and slight cross-beak deformities. The analysis prioritized several candidate genes and pathways that may be relevant to cross-beak deformity and phenotypic severity in Huiyang Bearded chickens.

Functional enrichment analysis of genes harboring candidate variants highlighted terms related to neural crest cell migration. Neural crest cells undergo epithelial-to-mesenchymal transition, delaminate from the dorsal neural tube, and migrate to multiple embryonic destinations, where they contribute to craniofacial structures and then further migrate from the neuroepithelium to different areas of the skull, forming various craniofacial structures. In the early stages of craniofacial embryogenesis, cranial neural cranial neural crest cells arising from the forebrain, midbrain, and hindbrain regions migrate to the ventral area of the head and fill the pharyngeal arches. Cranial neural crest cells contribute to craniofacial connective tissues, cartilage, bone, and other structures that support development of the upper and lower beaks and the anterior skull [10]. The enrichment of neural-crest-related annotations suggests that altered developmental processes involving neural crest cells may warrant investigation in cross-beak deformity. Future studies should investigate whether the prioritized variants influence neural crest cell migration or other craniofacial developmental processes.

*CDH11* and *DHFR* were prioritized as candidate genes because they harbored candidate variants and occupied connected positions in the exploratory PPI network. NC cells are multipotent embryonic cells that form craniofacial bone and cartilage. Previous developmental studies have suggested that *CDH11* may contribute to neural crest cell morphology and migration; however, its role in chicken cross-beak deformity remains to be experimentally evaluated. [21]. Folate deficiencies or mutations to major components can result in craniofacial defects [22, 23], which may be a consequence of neural tube closure and cranial neural crest defects [24, 25]. *DHFR* participates in folate metabolism by catalyzing the reduction of dihydrofolate to tetrahydrofolate, thereby supporting one-carbon metabolic processes required for nucleotide biosynthesis and cellular proliferation., which is required in the presumptive face during embryonic mouth and facial prominence specification [26]. Inhibiting *DHFR* during early orofacial development severely reduces later neural crest derivatives, such as jaw cartilage, as well as mesoderm derivatives, such as muscle [27]. Cross-beak deformity may arise from perturbations in craniofacial development; however, the developmental stages and molecular processes involved remain incompletely understood. Our findings nominate *CDH11* and *DHFR* as candidate genes for functional studies of cross-beak deformity.

Genes harboring candidate SNPs, including *NRXN3, NRXN1, CDH5,* and *SDC3*, were represented in the cell adhesion molecule pathway in the enrichment analysis. The genes *NRXN1* and *NRXN3* are transmembrane synaptic proteins of the central nervous system and are involved in cell adhesion, cell recognition, and blood vessel formation. Evidence from human craniofacial studies may suggest a potential developmental relevance of *NRXN3*; however, no direct evidence currently links *NRXN3* variation to cross-beak deformity in chickens [28]. The biological role of *CDH5* is to regulate the aggregation and separation of cells of the same type and to promote the formation of tissues and organs during embryogenesis [29]. Cadherin regulates cell‒cell adhesion and prevents neural crest cells from moving forward through contact inhibition of locomotion (*CIL*). Inhibiting the expression of focal adhesive junctions during cell‒cell contact indicates that *CDH5* plays an important role in early craniofacial development in chickens [30]. *SDC3* is a member of the transmembrane proteoglycan family (syndecans), which is closely related to osteogenic differentiation [31]. The planar cell polarity (*PCP*) pathway is known to interact with syndecan to control the direction and migration direction of cell protrusions, thereby affecting the migration of neural crest cells [32]. Given the reported roles of syndecans in cell migration and osteogenic processes, *SDC3* represents a candidate for future investigation in craniofacial development In addition, with respect to the GO gene enrichment results, the most significant pathway was the single-organ cell‒cell adhesion pathway, indicating the potential function of cell adhesion in the formation of cross-beaks.

Based on the different analyses of high and slight cross beak chickens, Comparative analysis between the severe and slight deformity groups identified candidate variants in *MRPL21, NSUN2, DDX55, GNB3*, and *NFKB2*; these genes were prioritized as hypotheses for future studies of phenotypic severity. The involvement of *MRPL21*, a member within the family of mitochondrial ribosomal proteins (MRPs), is well-documented in both cellular apoptosis and energy metabolism [33]. *NSUN2* is a key methylating molecule in RNA m5C modification, acting as a methyltransferase to transfer methyl groups to tRNA, mRNA, and non-coding cytosine residues on RNAs [34]. *DDX55*, a member of the DEAD-box RNA helicases, belongs to helicase superfamily, could induce cell cycle progression and EMT by activating β -catenin in a BRD4/PI3K/Akt/GSK-3 β-dependent manner [35]. G protein is an important component of a variety of transmembrane receptors involved in bone metabolism, and *GNB3* is the beta3-subunit of G protein. *GNB3* encodes a G-protein β subunit and may influence signaling pathways relevant to cellular responses; its potential relevance to cross-beak deformity remains unknown [36]. Osteoporosis is a common bone metabolic disease, and *NFKB2* has been proven to be a potential pathogenic gene for osteoporosis [37, 38]. Cross beak is a beak deformity. Notably, the occurrence of craniofacial development during embryonic development is one of the main reasons resulting in malformed beaks; therefore, genes related to craniofacial development may be associated with the formation of cross-beaks. Craniofacial development is related to the development of cartilage, bone matrix and blood vessels. Thus, we mainly focused on the genes with these functions in the difference analysis. The research analysis of this study showed that the *MRPL21, NSUN2, DDX55, GNB3, and NFKB2* genes were related to the degree of morphological abnormalities of beak deformities by enrichment in the focal adhesion, amyotrophic lateral sclerosis, primary bile acid biosynthesis signaling pathways, chemokine signaling pathway, c-type lectin receptor signaling pathway, and relaxin signaling pathway.

*BMP4*, which has previously been proposed as a candidate gene based on expression and physiological observations [19, 39], was not prioritized in the present variant-based analysis. This discrepancy may reflect differences between expression-based and variant-based approaches, limited sample size, phenotypic heterogeneity, or incomplete coverage of relevant regulatory variation. Future studies integrating independent-cohort genotyping, gene-expression profiling, and functional assays are needed to evaluate BMP4 together with the candidate genes identified here.

Sanger sequencing confirmed technical concordance for selected variants detected by whole-genome resequencing; however, it did not provide independent replication of genotype–phenotype associations. Validation in larger, independent cohorts that include both affected and normal-beaked chickens will be required to estimate allele frequencies, evaluate genotype–phenotype relationships, and assess the reproducibility of the prioritized variants.

This study has several limitations. In particular, the small sample size, including the limited number of normal-beaked chickens, precluded genome-wide association testing and robust inference regarding variant effects. The results should therefore be interpreted as exploratory comparative findings rather than evidence of causal genetic determinants of cross-beak deformity. Future studies should combine larger and more balanced populations, pedigree information, population-structure assessment, gene-expression profiling, and functional experiments to clarify the biological relevance of the candidate variants and genes identified here..

## DECLARATIONS

### Author contributions

The experiment was mainly conceived and designed by Hua Li and Hui Yu; the experiment was performed by Fei Ye, Haiyi Yu, Yuyu Hong, Haiquan Zhao and Huiming Kang; experimental data were collected and analyzed by Yuyu Hong; the manuscript was mainly written by Fei Ye, and assistance and critical revisions to the draft were provided by Yuyu Hong, Haiyi Yu, and Hua Li; Hua Li resourced the project. All the authors read and approved the final manuscript.

### Availability of data and materials

All raw resequencing data used in this study have been deposited into the NCBI BioProject database under accession PRJNA839907.

### Consent for publication

Not applicable.

## Acknowledgment

Not applicable.

## Supporting information

Supplementary file 1-Bioinformatics analysis of SNP variant genes of cross-beak group and normal group.

Supplementary file 2-The results of indel annoted genes of cross-beak group and normal group.

Supplementary file 3-GO analysis of indel variant genes of cross-beak group and normal group.

Supplementary file 4-KEGG analysis of indel variant genes of cross-beak group and normal group.

Supplementary file 5-The string interactions of indel annoted genes of cross-beak group and normal group.

Supplementary file 6-The string_node_degrees of indel annoted genes of cross-beak group and normal group.

Supplementary file 7- The GO enrichment of SNP annotated genes of high and slight cross-beak chickens.

Supplementary file 8-The KEGG enrichment of indel annotated genes of high and slight cross-beak chickens.

Supplementary file 9- The GO enrichment of indel annotated genes of high and slight cross-beak chickens.

